# Metformin Suppresses SARS-CoV-2 in Cell Culture

**DOI:** 10.1101/2021.11.18.469078

**Authors:** Haripriya Parthasarathy, Dixit Tandel, Abdul Hamid Siddiqui, Krishnan H. Harshan

**Affiliations:** CSIR-Centre for Cellular and Molecular Biology, Hyderabad, India 500007; Academy for Scientific and Innovative Research (AcSIR), Ghaziabad-201002, India

**Keywords:** SARS-CoV-2, COVID-19, antiviral activity, Metformin, AMPK, Diabetes

## Abstract

Comorbidities such as diabetes worsen COVID-19 severity and recovery. Metformin, a first-line medication for type 2 diabetes, has antiviral properties and certain studies have also indicated its prognostic potential in COVID-19. Here, we report that metformin significantly inhibits SARS-CoV-2 growth in cell culture models. First, a steady increase in AMPK phosphorylation was detected as infection progressed, suggesting its important role during viral infection. Activation of AMPK in Calu3 and Caco2 cell lines using metformin revealed that metformin suppresses SARS-CoV-2 infectious titers up to 99%, in both naïve as well as infected cells. TCID50 values from dose-variation studies in infected cells were found to be 0.8 and 3.5 mM in Calu3 and Caco2 cells, respectively. Role of AMPK in metformin’s antiviral suppression was further confirmed using other pharmacological compounds, AICAR and Compound C. Collectively, our study demonstrates that metformin is effective in limiting the replication of SARS-CoV-2 in cell culture and thus possibly could offer double benefits s diabetic COVID-19 patients by lowering both blood glucose levels and viral load.

## INTRODUCTION

COVID-19 prevails and rages on in several parts of the world. While vaccination drives continue to ensure its spread is limited, newer worrying aspects of long-COVID are a cause for concern. Patients with comorbidities such as cancer, auto-immune diseases, cardiovascular conditions, and diabetes, are still highly susceptible to contracting the disease (1,2). Particularly vulnerable to COVID-19 are patients with comorbidities such as cancer, auto-immune diseases, cardiovascular conditions, and diabetes. The hospitalization rate for patients with comorbidities who contracted COVID-19 was significantly higher during the first wave of the pandemic, associated with poor prognosis (3). Growing evidence of long-COVID indicates that long-term effects of the illness along with a possibility of latency of SARS-CoV-2 can drastically affect the complete recovery of COVID-19 patients (4,5). Evidences of SARS-CoV-2 for prolonged periods have been observed in lung (6), and intestinal epithelium (7). The prospect of repurposing drugs to combat prolonged presence of the virus is of utmost importance to manage disease outcomes.

Type 2 diabetes, one of the most common metabolic disorders, is universally treated using insulin and a number of other drugs, chiefly metformin (8). Metformin is a biguanide compound, used as first-line antidiabetic medication worldwide. It acts primarily by increasing glucose intake and limiting gluconeogenesis in the liver, and its action is mediated in part by the energy-sensing kinase, 5’-AMP-activated protein kinase (AMPK) (9). Metformin inhibits Complex I of the electron transport chain and suppresses ATP synthesis, which triggers AMPK activation. This results in a cascade of events that decreases anabolic processes and initiates macromolecular breakdown to reinstate homeostasis. AMPK is a trimeric complex where the α subunit harbors the kinase and activating domains. The regulatory subunit γ is where AMP or ATP binds, thereby determining the allosteric regulation of AMPK (10).

Competitive binding of AMP or ATP to one of four binding domains on the γ subunit is believed to cause a conformational change that makes the kinase domain on the α subunit more accessible for phosphorylation, although this notion has been debated (10,11).

In the past year, several reports have debated the clinical use of metformin in COVID-19 (12–15). Many case studies on metformin treatment report a decrease in hospital mortality rates for patients that were on metformin prior to admission (13,16,17). Reports suggest that high glucose levels are associated with poorer prognosis and well-controlled glucose levels were indicative of lesser complications (16). Metformin has also been reported to show antiviral activity against other viruses (18). In this study, we aimed to investigate the effect of metformin on SARS-CoV-2, and identify the effects of AMPK perturbation on infection. With recent reports having clarified that COVID-19 is not solely a respiratory illness with evidence of SARS-CoV-2 found in feces of patients showing successful replication in enterocytes (19,20) we hence used cell lines Calu3 and Caco2, a lung carcinoma and a colorectal carcinoma cell line, respectively. Treatment of cells with metformin prior to infection substantially lowered the viral titer. Additionally, metformin strongly restricted SARS-CoV-2 levels in previously infected cells in a dose-dependent manner. Pharmacological activation of AMPK through AICAR suppressed viral infection while its inhibition using Compound C promoted it, confirming the effect of metformin is through AMPK. Our results support the promising use of metformin as a therapeutic drug in COVID-19.

## MATERIALS AND METHODS

### Cell culture and reagents

Caco2, Calu3, and Vero cells were grown in DMEM supplemented with FBS, and Pen Strep, at 37°C and 5% CO_2_. Antibodies against AMPK and phospho-AMPK (T172) were procured from CST. GAPDH, β-tubulin, and Nucleocapsid antibodies were from ThermoFisher Scientific. HRP-conjugated secondary antibodies were purchased from Jackson ImmunoResearch. Metformin, AICAR, and Compound C were procured from Merck Millipore.

### Infections and treatments

All experiments involving virus culture were carried out in the biosafety level-3 laboratory at the Centre for Cellular and Molecular Biology (CCMB). SARS-CoV-2 strain B.1.1.8 (TG-CCMB-L1021/2020 isolate) was used for all experiments at 1 MOI (21). Cells were grown to 80% confluency and treated as described. For pre-infection treatments, cells were subjected to 10 mM metformin or 1 mM AICAR for 24 h, followed by infection with SARS-CoV-2 in serum-free medium (SFM) for 3 h in the presence of the respective compound. The inoculum was subsequently replaced with complete medium containing the compound and the cells were harvested at 24 h post-infection (hpi). In post-infection mode of metformin treatment, the cells were first infected for 3 h after which the inoculum was replaced by media containing 10 mM metformin and further incubated until 24 hpi. Dose-dependent effect of metformin was studied by subjecting cells to varying doses of metformin (5, 10, 20, and 40 mM in Caco2, and 1, 2, 5, 10, and 20 mM in Calu3 cells) similar to the post-infection treatment regimen mentioned above. Compound C treatment was carried out by infecting cells for 3 h at 1 MOI, complete media for 21 h, and then an additional 24 h with 10 µM Compound C. For all treatments, media supernatant was collected to measure extracellular viral RNA as well as infectious viral titers, and cells were processed for immunoblotting.

### Virus quantification and titration

RNA from viral supernatants was isolated using Nucleospin Viral RNA isolation kit (Macherey-Nagel GmbH & Co. KG). qRT-PCR was carried out using nCOV-19 RT-PCR detection kit from Q-line Molecular to quantify SARS-CoV-2 RNA following manufacturer’s protocol on Roche LightCycler 480.

Infectious titers of the supernatants were calculated using plaque forming assay (PFU/mL) as mentioned previously (21). Briefly, the supernatant was serially diluted in SFM and added to a confluent monolayer of Vero cells for infection for 3 h. The medium was then replaced with a 1:1 mixture of agarose: 2 × DMEM (1% low-melting agarose (LMA) containing a final concentration of 5% FBS and 1 × Pen-Strep). Six days post-infection, cells were fixed in 4% formaldehyde prepared in 1 × PBS and subsequently washed and stained with 0.1% crystal violet to count the plaques.

### Immunoblotting

Cell pellets were lysed in an NP-40 lysis buffer as described earlier (21). Protein quantification was done using BCA method (G Biosciences). Lysates were then mixed with 6 × Laemmli buffer, and equal amounts of protein were run on SDS-PAGE, followed by transfer onto PVDF membrane. Blots were blocked in 5% BSA and incubated with specific primary antibodies at 4°C overnight. Incubation with HRP-conjugated secondary antibodies was done for 1 hour and the blots were developed on a BioRad Chemidoc MP system using ECL reagents (ThermoFisher and G Biosciences). Quantification was performed using ImageJ (22).

### Statistical analysis

All experiments were performed in triplicate to calculate mean ± SEM. Statistical significance was calculated using two-tailed, unpaired Student’s *t*-test and *p* values are represented as *, **, ***, indicating *p* ≤ 0.05, 0.005, and 0.0005, respectively.

IC50 and IC90 were calculated from qRT-PCR data, while TCID50 and TCID90 were calculated from PFU data from metformin titration experiments.

## RESULTS

### SARS-CoV-2 infection causes long-term phosphorylation of AMPK

We tracked the activity and levels of AMPK during SARS-CoV-2 infection over a 96-h time course beginning from 1 hour. Caco2 cells were infected with 1 MOI of SARS-CoV-2 for 1, 2, 6, 12, 24, 48, 72, and 96 h time. Though no significant change in the AMPK phosphorylation was detected until 48 h post-infection (hpi), marked increase in phosphorylation was evident from 48 hpi that further strengthened until 96 hpi, despite a drop in the abundance of the protein (Figure 1 A and B). These results indicated a major metabolic reprogramming, resulting in AMPK phosphorylation occurring after 24 hpi. Interestingly, AMPK phosphorylation coincided with the accumulation of viral proteins (Figure 1A).

**Figure 1.**
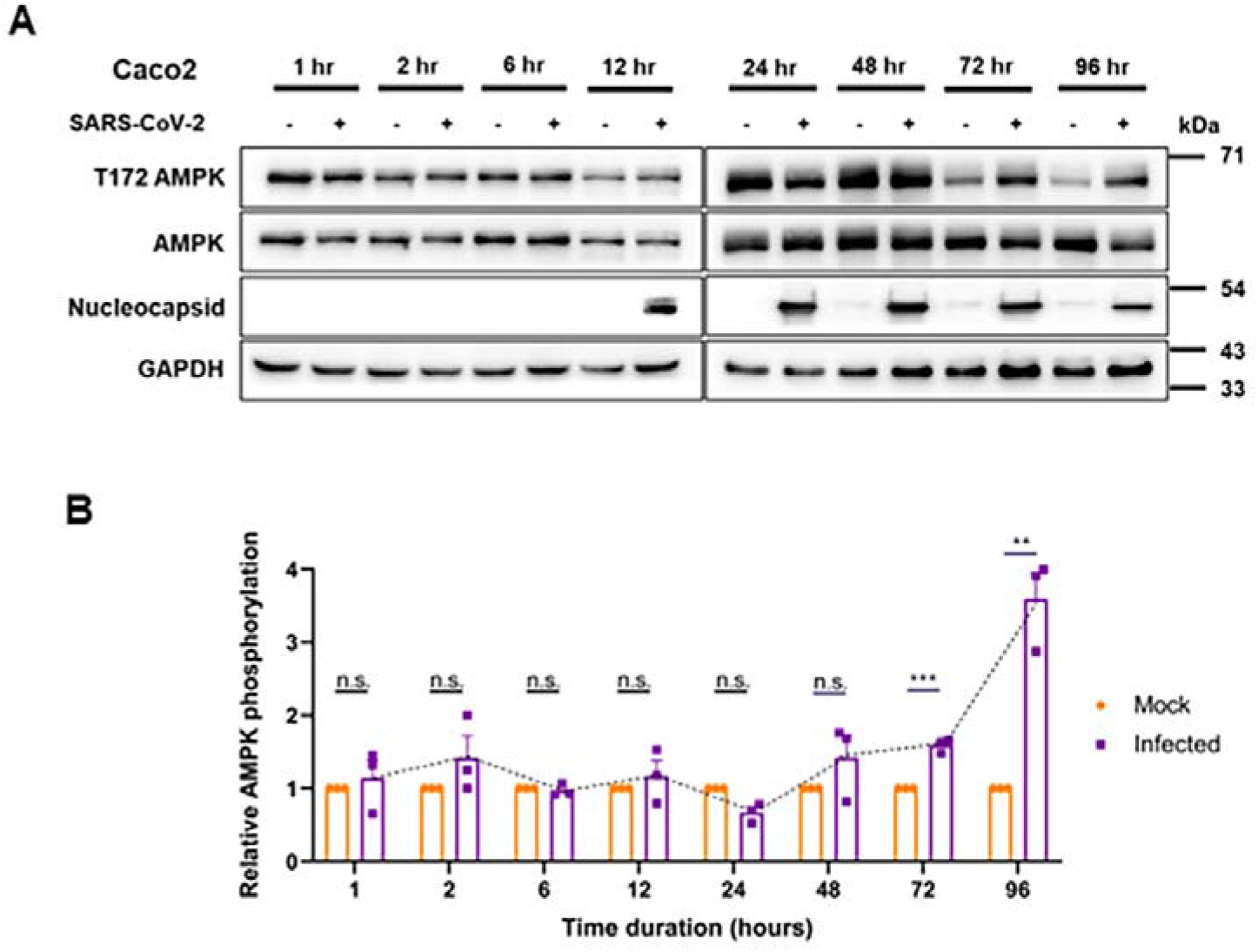
SARS-COV-2 infection induces AMPK phosphorylation. **(A)** Immunoblots analyzing the phosphorylation of AMPK in SARS-CoV-2 infected Caco2 cells for early (1, 2, 6, and 12 h) and late (24, 48, 72, and 96 h) time points. Cells infected with 1 MOI of SARS-CoV-2 were harvested at various time intervals post-infection and analyzed by immunoblotting. **(B)** Quantitative representation of AMPK phosphorylation from three independent replicates. Densitometric values of phospho-AMPK bands were normalized against those of total-AMPK expression and GAPDH belonging to the corresponding samples and the values were plotted graphically.

### Metformin protects cells from SARS-CoV-2 infection

We investigated the role of AMPK activation during SARS-CoV-2 infection as previous reports have clearly established the roles played by this molecule on the outcome of viral infections (23). One compound of clinical relevance known to activate AMPK is metformin. With the emergence of reports suggesting a possible beneficial role of metformin in COVID-19 patients (12,13), we used this compound to both activate AMPK, as well as study it from a drug-repurposing point of view. In this study, we used Calu3, a respiratory epithelial cell line, and Caco2, a gut epithelial cell line, allowing us to study a larger impact of metformin and extrapolate its results in cells from different organs of relevance. Calu3 cells were pre-treated with 10 mM concentration of metformin for 24 h after which they were infected with 1 MOI of SARS-CoV-2 for 3 h in presence of metformin. Metformin at 10 mM is widely used in studies, is physiologically relevant and well reported to induce AMPK activity (24– 26). Subsequently, the inoculum was replaced with growth medium containing metformin and incubated until 24 hpi (Figure 2A). Increased phosphorylation of AMPK in the drug-treated cells confirmed the effect of metformin (Figures 2 B and C). Metformin treatment resulted in nearly 90% drop in the viral RNA (Figure 2D), and nearly two-log drop in the infectious viral titers (Figure 2E) indicating that metformin treatment is protective against SARS-CoV-2 infection. Viral protein levels also decreased by 50%, suggesting a clear suppression of the viral life cycle by metformin (Figure 2B). Metformin treatment performed in Caco2 cells corroborated the results from Calu3, confirming that the drug has strong protective effects against SARS-CoV-2 infection in multiple cell lines (Figures 2 F-I).

**Figure 2.**
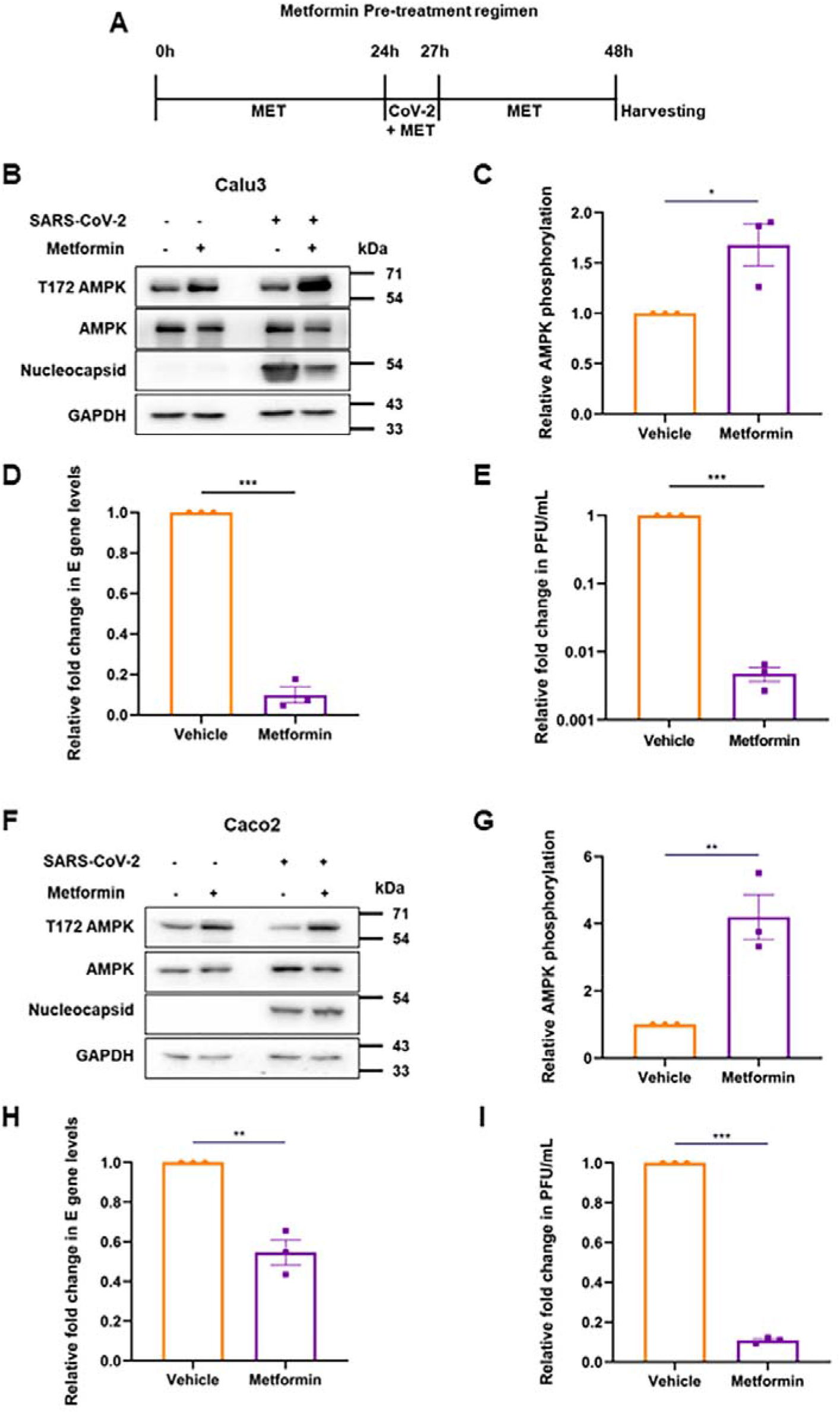
Metformin pre-treatment inhibits SARS-CoV-2. **(A)** Schematic of the experimental set up for pre-treatment. Cells were infected with 1 MOI SARS-CoV-2 24 h after 10 mM metformin treatment. The vehicle control cells were also infected with the virus in parallel. **(B)** Immunoblots from Calu3 cells confirming AMPK phosphorylation by metformin or vehicle treatment, and SARS-CoV-2 infection. **(C)** Densitometric quantification of AMPK phosphorylation in infected cells. **(D)** SARS-CoV-2 RNA levels in the supernatant of metformin treated samples, measured by qRT-PCR of E gene. Relative fold change in the E levels between metformin and vehicle treated samples is depicted. **(E)** Relative fold change in the infectious viral titers of SARS-CoV-2 in metformin treated samples compared against that treated with vehicle, represented as fold change in PFU/mL. Similar experiments were carried out in Caco2 cells, with immunoblots representing AMPK phosphorylation **(F)**, its quantification **(G)**, viral RNA **(H)** and infectious units **(I)**. Graphs indicate mean ± SEM, indicating the three biological replicates.

### Metformin treatment post-infection causes more profound restriction of SARS-CoV-2

We next tested the effect of metformin in cells previously infected with SARS-CoV-2 to extrapolate its impact on the infected patients. Cells infected with 1 MOI of SARS-CoV-2 for 3 h were subsequently treated with 10 mM metformin until harvested at 24 hpi (Figure 3 A). Metformin caused higher AMPK phosphorylation in both Calu3 and Caco2 cultures infected with SARS-CoV-2 (Figures 3 B and C, F and G). Post-infection treatment with metformin resulted in a significant decrease in viral RNA and almost one-log drop in infectious titer in the supernatant of Calu3 cells with reduction in nucleocapsid expression as well (Figures 3 B, D, and E). Similar results were observed in Caco2 cells, including a two-log reduction in viral titers (Figure 3 G-K), further confirming the profound restrictive effect of the drug on SARS-CoV-2. These results collectively indicate that metformin offers strong protection to both naïve as well as infected cells.

**Figure 3.**
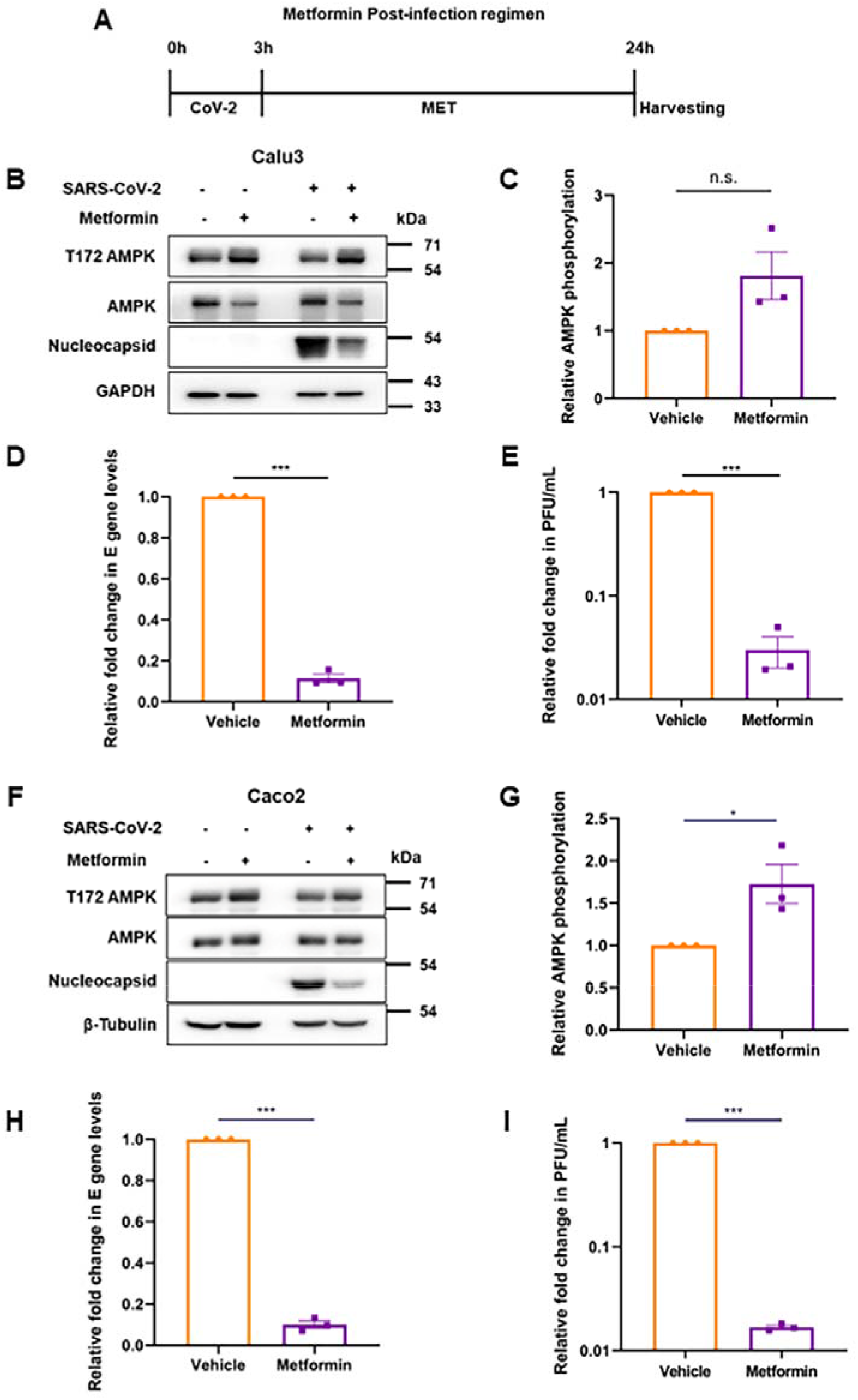
Metformin suppresses SARS-CoV-2 replication in the infected cells. **(A)** Schematic of the post-infection treatment by metformin. Cells were infected with 1 MOI SARS-CoV-2 for 3 hours, followed by 21 h of metformin treatment before harvesting at 24 hpi. Vehicle treated cells were infected in parallel. **(B)** Confirmation of SARS-CoV-2 infection in Calu3 cells and AMPK phosphorylation by immunoblotting. **(C)** Densitometric analysis of AMPK phosphorylation in the treated, infected cells. **(D)** Relative fold change in SARS-CoV-2 E gene measured by qRT-PCR in the supernatants of metformin treated cells compared with the those treated with vehicle. **(E)** Relative SARS-CoV-2 infectious titers of the supernatant from samples treated with metformin represented as fold change in PFU/mL. Similar experiments were performed in Caco2 cells. Confirmatory immunoblots **(F)**, quantification of AMPK phosphorylation **(G)**, RNA levels in the supernatant **(H)**, and infectious titers from vehicle and treated cells **(I)**, are represented. Graphs indicate mean ± SEM, indicating the three biological replicates.

### Metformin treatment post-infection causes a dose-dependent suppression of SARS-CoV-2

Metformin dose administered to diabetic individuals varies widely depending on a number of clinical parameters including fasting plasma glucose levels and HbA1c (27). Hence, based on our hypothesis, it is possible that individuals prescribed with different metformin doses are affected differently when infected with SARS-CoV-2. To assess whether *in vitro* administration of different doses of metformin affected the virus differently, we performed a dose-variation analysis. A concentration-dependent increase in AMPK phosphorylation was evident in SARS-CoV-2 infected cells from 1-20 mM metformin concentrations in Calu3 cells (Figures 4 A and B). A gradual and dose-dependent decrease in N levels was also evident (Figures 4 A and C). The drop in viral RNA levels was more dramatic with a very significant drop detected even at 1 mM concentration, that stabilized between 2-20 mM concentration (Figure 4D). Drop in viral titers was even more profound, with viral titers sharply decreasing beyond 2 mM metformin (Figure 4E). The IC50 calculated using RNA values was 0.8 mM (Figure 4F) with TCID50/90 values from PFU being 2.0 and 4.9 mM, respectively (Figure 4G). A comparable decrease in N, viral RNA, and infectious titers was observed in Caco2 cultures with increasing metformin dose (Figure 4 H-L). IC50 value was calculated to be 2.9 mM with a corresponding TCID50 value of 3.5 mM (Figure 4 M and N). Together, these results unambiguously demonstrate a potent anti-SARS-CoV-2 effect of metformin.

**Figure 4.**
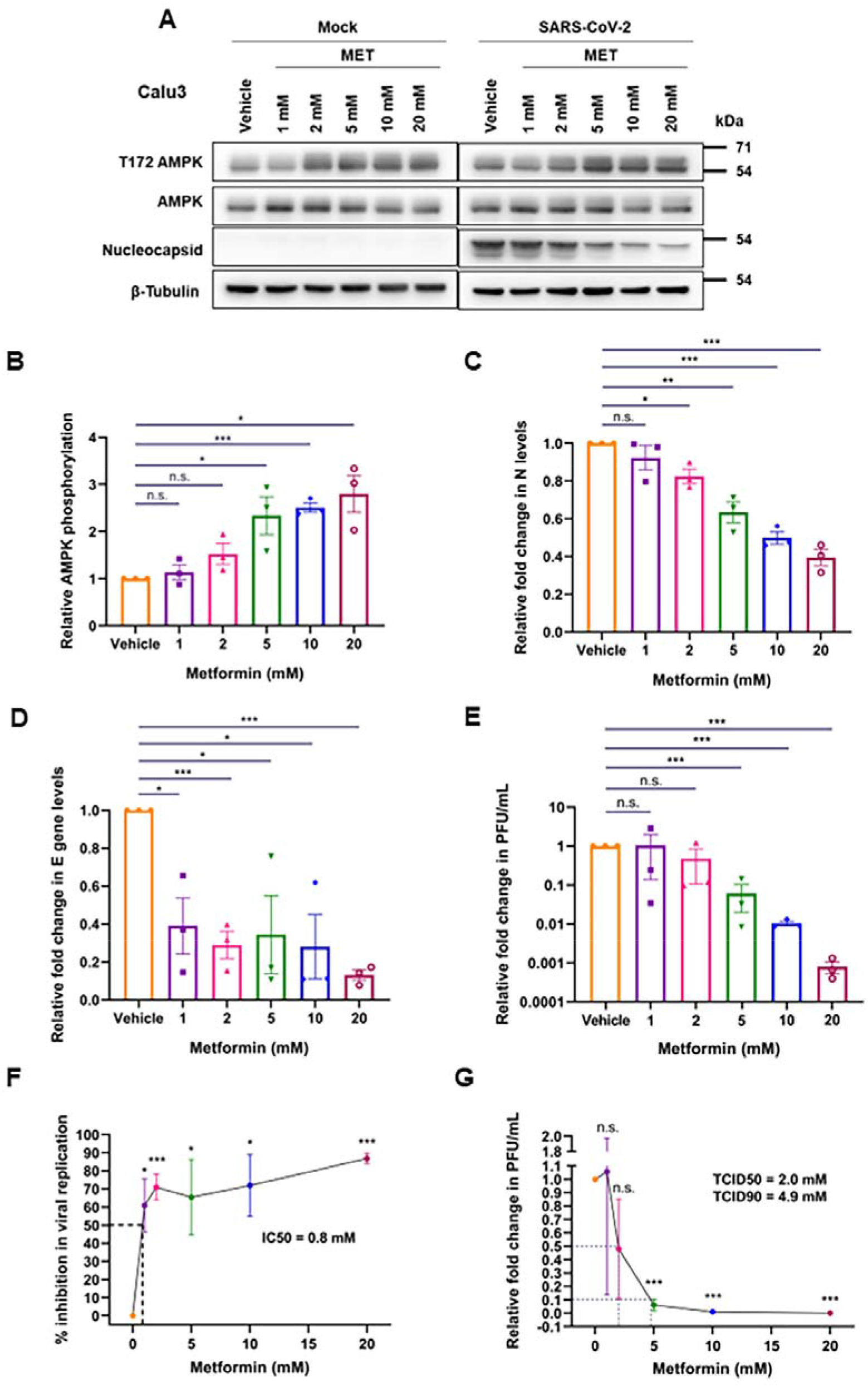
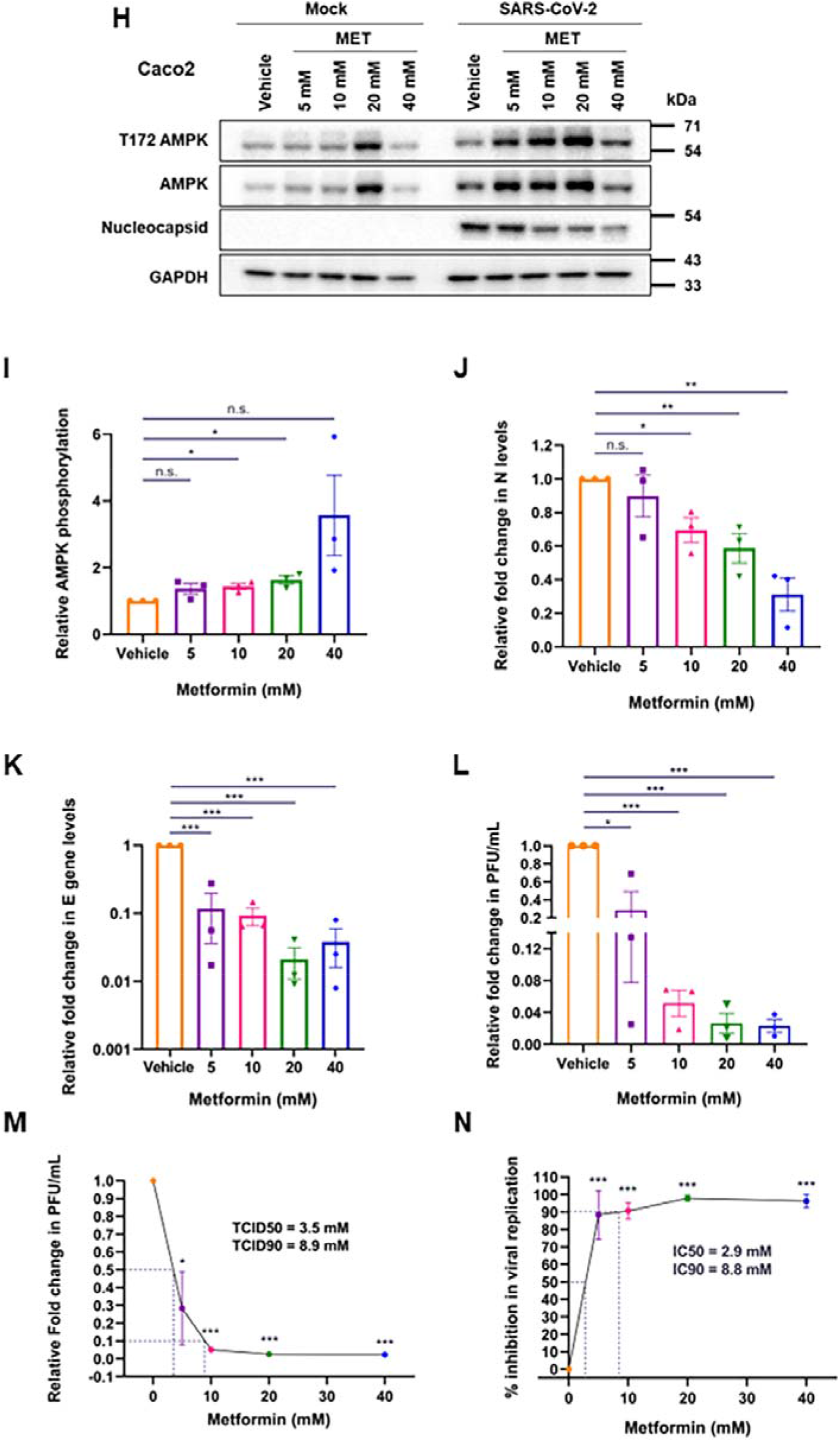
Metformin treatment inhibits SARS-CoV-2 replication in a dose-dependent manner. **(A)** Immunoblots confirming the infection and AMPK phosphorylation following the treatment of SARS-CoV-2 infected Calu3 cells with metformin at the doses described above the panel. **(B)** Relative AMPK phosphorylation in the samples treated with metformin. The graph was generated from the densitometric analysis of the immunoblots. **(C)** Relative abundance of N quantified from the immunoblots from the panel (A). **(D)** Relative fold change in SARS-CoV-2 E gene measured by qRT-PCR in the supernatants of metformin treated cells compared with the those treated with vehicle. **(E)** Relative SARS-CoV-2 infectious titers of the supernatant from samples treated with metformin represented as fold change in PFU/mL. **(F)** Calculation of IC50 and IC90 for metformin on SARS-CoV-2 replication measured by plotting E gene levels present in the supernatants of the samples treated with the respective concentrations of metformin. **(G)** Measurement of TCID50 and TCID90 for metformin on SARS-CoV-2 infectious virus particle production. PFU/mL data for the individual samples treated with the specific concentrations were plotted in the graph to calculate the respective values. **(H)** Immunoblots confirming infection and AMPK phosphorylation in SARS-CoV-2 infected Caco2 cells treated with metformin at the doses mentioned. **(I)** Relative AMPK phosphorylation in the metformin treated samples. **(J)** Relative abundance of N quantified from the immunoblots from (H). **(K)** Relative fold change in SARS-CoV-2 E gene measured by qRT-PCR in the supernatants of metformin treated cells compared with vehicle control. **(L)** Relative SARS-CoV-2 infectious titers from varying metformin treatments **(M)** IC50 and IC90 from SARS-CoV-2 E gene levels present in the supernatants. **(N)** TCID50 and TCID90 from data depicted in (L). Graphs indicate mean ± SEM, indicating the three biological replicates.

### AMPK activation restricts SARS-CoV-2

We then sought to understand the mechanism by which metformin may be carrying out its suppressive effects. Since metformin’s mode of action through AMPK is well established, we wanted to evaluate whether AMPK mediated this effect. To verify the same, we used pharmacological compounds that would activate or inhibit AMPK activity. AICAR, an allosteric activator, directly activates AMPK by binding to the γ subunit, mimicking AMP-binding, thereby activating downstream signaling. Cells were pre-treated with 1 mM of AICAR, a well-established concentration (28,29). for 24 h as in the case of metformin (Figure 5A). AICAR treatment activated AMPK (Figure 5B and C) with a marginal change in nucleocapsid (Figure 5A), while significantly lowering the infectious viral titer of SARS-CoV-2 (Figure 5D) by almost one-log, as demonstrated previously with metformin (Figure 2I). These results demonstrate that AMPK activation is certainly beneficial to the host cells by significantly limiting the viral titers. We further confirmed this effect by inhibiting AMPK by using Compound C (CC), a small molecule ATP-competitive inhibitor, during SARS-CoV-2 infection. Since AMPK phosphorylation peaked beyond 24 h in the time-course study (Figure 1A), we decided to infect the cells first to activate AMPK and then inhibited AMPK with CC. Cells infected at 1 MOI were treated with 10 µM CC at 24 hpi and incubated for an additional 24 h (Figure 5E), for a total of 48 hours of infection. CC is used at this concentration widely (9,30). Though CC treatment caused a visible drop in AMPK phosphorylation in mock cells, there was no apparent decrease observed in infected cells, indicating that the virus-induced AMPK activation overrides CC inhibition (Figure 5 F and G). As anticipated, CC treatment resulted in over four-fold higher viral titers in the supernatants as against the control sample (Figure 5H). These results confirm that AMPK coordinates strong antiviral measures in SARS-CoV-2 infected cells. Thus, activation of AMPK during the infection is protective against SARS-CoV-2 infection.

**Figure 5.**
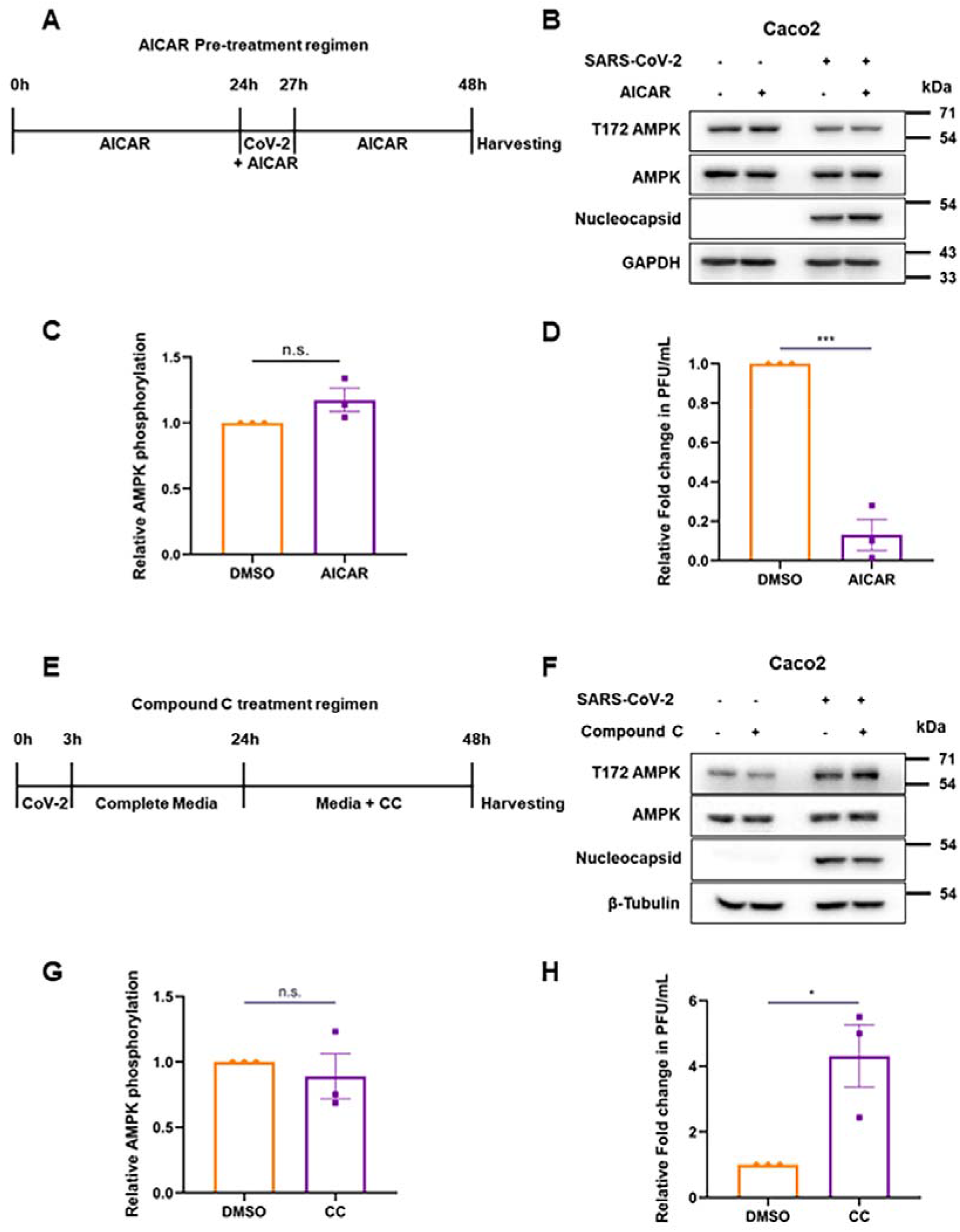
AMPK activation is beneficial to the host against SARS-CoV-2. **(A)** Schematic of the treatment of Caco2 cells with AICAR and infection by SARS-CoV-2. **(B)** Immunoblots depicting AMPK phosphorylation and levels, and confirming infection. **(C)** Relative AMPK phosphorylation from AICAR-treated, infected cells. **(D)** Relative infectious titers of SARS-CoV-2 in samples that underwent pre-treatment with AICAR, as against the vehicle. **(E)** Schematic of the treatment of SARS-CoV-2 infected Caco2 cells with CC. **(F)** Immunoblot confirmation of the infection and inhibition of AMPK activity. **(G)** Quantification of AMPK phosphorylation in CC treated, infected samples. **(H)** Relative infectious titers of SARS-CoV-2 in samples that underwent CC-treatment, as against the vehicle, DMSO. Graphs indicate mean ± SEM, indicating the three biological replicates.

## DISCUSSION

The anti-diabetic drug, metformin, has been projected to influence the prognosis of COVID-19 patients. As patients with comorbidities fared worse when infected with SARS-CoV-2, management of an ongoing illness alongside COVID-19 treatment became paramount. Some studies early during the pandemic identified a positive correlation between improved glucose levels in diabetic patients on metformin and better clinical outcome (16,31–33). A number of reports proposed metformin as a possible “miracle or menace” in COVID-afflicted patients, based on retrospective data from hospitalisations, comparing the length of hospitalisation, severity of symptoms, or mortality (17,34). In this study, we demonstrate that metformin profoundly lowers SARS-CoV-2 infectivity. While extrapolating these results to a clinical setup may not be appropriate, our data indicate that metformin can be effective not only as a treatment option, but as a prophylactic agent as well. With this data, we suggest that treatment of patients with metformin prior to infection with SARS-CoV-2 may have assisted in decreasing their symptoms of COVID-19. Our results also suggest that metformin could be beneficial in non-diabetic, COVID-19 patients and expand the scope of its coverage. In summary, this data lies in agreement with the numerous case studies published during the pandemic that suggested an antiviral role for this known anti-diabetic drug.

Metformin plays a major role in modulating lipid metabolism, but the mechanisms are multi-dimensional. Metformin is a soluble compound that interferes with Complex I of the electron transport chain, and the decrease in ATP production causes AMPK activation. It also decreases hepatic lipids, increases skeletal muscle uptake of glucose, and in parallel helps in decreasing circulating lipids that can eventually increase cardiovascular risk especially in diabetic adults who are obese (35,36).

RNA viruses in particular are known to modulate lipogenesis to steer cells to produce more vesicles to aid replication, as well as packaging and release (37,38). Certain viral infections trigger induction of lipogenic genes through activating factors such as SREBPs and PPARs, resulting in accumulation of lipid droplets (39,40). These facilitate the establishment of key sites of replication and assembly, which has been documented for several viruses including flaviviruses (41–44) and others (45–47). This can also act as a double-edged sword as a number of antiviral molecules also assemble on lipid vesicles. Activation of interferon pathways, particularly types I and III, results in the production of interferon-stimulated genes (ISGs), some of which are reported to load onto lipid droplets, especially viperin and IFIT-1 (48). A recent report on SARS-CoV-2 highlighted the possible role that lipid droplets play in its infection (49). Not only did they observe higher colocalization of viral RNA with the lipid droplets, they also demonstrated that inhibition of its formation decreased viral load as well as pro-inflammatory cytokines and apoptosis markers. These results in conjunction with ours indicate that the anti-viral effect of metformin is probably a resultant of altered lipid metabolism.

Certain RNA viruses have been found to remain in the host long after onset of symptoms. Some studies have observed the presence of SARS-CoV-2 RNA several weeks after symptoms develop, leading to prolonged illness (6). Presence of viral RNA for extended periods indicates the possibility of latency in COVID-19, though not proven yet. These factors are predicted to play a key role in long COVID-symptoms observed in a large number of individuals that have been infected (50).

From the results of this study, it is likely that the use of metformin in such cases could be an interesting proposition to combat latent COVID-19.

We conducted studies in Caco2 and Calu3, two cell lines with pertinent tissue origin to our study. Metformin is absorbed in the intestine, which makes Caco2 an appropriate choice, in addition to its permissivity to SARS-CoV-2. The use of metformin in lung cells, an unusual choice with respect to metformin use, was in order to study the suppressive effects of metformin in SARS-CoV-2 permissive cells. We observed metformin’s potent and impressive effect on lung as well as gut cells, suggesting the possibility that other tissues where SARS-CoV-2 may infect would also respond to the drug similarly (51–54).

We speculate that the loss in infectivity of SARS-CoV-2 by metformin could also be an outcome of altered lipid metabolism mediated by AMPK. AMPK affects cellular lipid levels through a number of its substrates, such as ACC and SREBP1. AMPK also regulates macromolecular metabolism, mitochondrial homeostasis, autophagy as well as apoptosis. As its role is multifaceted and vital for maintaining energy levels, it has been reported to play key roles in many virus infections. Multiple reports show that AMPK activation can be either detrimental or beneficial for virus survival and propagation (23). Our results using AICAR and CC in SARS-CoV-2 infection implies an unfavorable/antiviral environment for the virus when AMPK is activated. In this context, it is interesting to note that N protein was unaffected during pharmacological activation of AMPK unlike the viral RNA and infectious titer, indicating that viral protein translation may not be inhibited during the treatments.

AICAR is an AMP analog that when phosphorylated, binds to the AMPK γ subunit. Accumulation of AICAR in the cytosol mimics increased AMP levels, skewing the AMP/ATP ratio thereby activating AMPK (11). Although a more specific activator of AMPK than metformin, AICAR does not necessarily have therapeutic potential, and hence we didn’t replicate all metformin experiments with AICAR. Compound C, or Dorsomorphin, is a largely-selective small molecule AMPK inhibitor that was picked up from a kinase inhibitor screen, for delineating the mechanism of action of metformin (9). Since we observed AMPK activation after 24 hours of infection, we wanted to inhibit AMPK when it is active. In our study, AMPK activation using AICAR reiterated the results of metformin albeit to a lower extent, while Compound C exhibited the opposite effect to metformin. These results suggest that although AMPK plays a pertinent role in bringing about the effects of metformin, there are other pathways involved in inhibiting SARS-CoV-2.

## CONCLUSION

Metformin suppresses SARS-CoV-2 *in vitro* and this data indicates both therapeutic and prophylactic potential for metformin in the management of COVID-19. AICAR and Compound C results explain AMPK’s partial role in mediating metformin’s antiviral effect.

## List of abbreviations

AMPK: AMP-activated protein kinase
ACC: Acetyl CoA carboxylase
AICAR: 5-Aminoimidazole-4-carboxamide ribonucleotide
CC: Compound C
COVID-19: Coronavirus disease 19
hpi: hours post infection
IFIT1: Interferon-induced protein with tetratricopeptide repeats 1
MOI: Multiplicity of infection
PPARs: peroxisome proliferator-activated receptors
SARS-CoV-2: Severe acute respiratory syndrome coronavirus 2
SREBPs: Sterol regulatory element-binding proteins
TCID50/90: 50%/ 90% Tissue culture infective dose

## Institutional biosafety

Institutional biosafety clearance was obtained for the experiments pertaining to SARS-CoV-2.

## Consent for publication

All authors agree to publish the article.

## Author contributions

H.P. performed treatments, infections, quantification and immunoblotting. D.N. performed qRT-PCR experiments. A.H.S. performed immunoblotting. H.P. and K.H.H. conceptualized the study and wrote the manuscript.

## Acknowledgement

We thank Divya Gupta, Vishal Sah, Sai Poojitha, and Prangya Paramita Sahoo for their help with generation of virus and for conducting experiments. We specially thank Mohan Singh Moodu and Amit Kumar for their assistance with logistics.

## REFERENCES

1. Gómez CE, Perdiguero B, Esteban M. Emerging SARS-CoV-2 Variants and Impact in Global Vaccination Programs against SARS-CoV-2/COVID-19. Vaccines [Internet]. 2021 Mar 1;9(3):1–13. Available from: /pmc/articles/PMC7999234/

2. Antonelli M, Pujol JC, Spector TD, Ourselin S, Steves CJ. Risk of long COVID associated with delta versus omicron variants of SARS-CoV-2. Lancet [Internet]. 2022 Jun 18;399(10343):2263–4. Available from: http://www.thelancet.com/article/S0140673622009412/fulltext

3. Sanyaolu A, Okorie C, Marinkovic A, Patidar R, Younis K, Desai P, et al. Comorbidity and its Impact on Patients with COVID-19. Sn Compr Clin Med [Internet]. 2020 Aug ;2(8):1. Available from: /pmc/articles/PMC7314621/

4. Pietsch H, Escher F, Aleshcheva G, Baumeier C, Morawietz L, Elsaesser A, et al. Proof of SARS-CoV-2 genomes in endomyocardial biopsy with latency after acute infection. Int J Infect Dis. 2021 Jan 1;102:70–2.

5. Davis HE, Assaf GS, McCorkell L, Wei H, Low RJ, Re’em Y, et al. Characterizing long COVID in an international cohort: 7 months of symptoms and their impact. EClinicalMedicine. 2021 Aug 1;38:101019.

6. Baang JH, Smith C, Mirabelli C, Valesano AL, Manthei DM, Bachman MA, et al. Prolonged Severe Acute Respiratory Syndrome Coronavirus 2 Replication in an Immunocompromised Patient. J Infect Dis [Internet]. 2021 Jan 1;223(1):23–7. Available from: https://pubmed.ncbi.nlm.nih.gov/33089317/

7. Natarajan A, Zlitni S, Brooks EF, Vance SE, Dahlen A, Hedlin H, et al. Gastrointestinal symptoms and fecal shedding of SARS-CoV-2 RNA suggest prolonged gastrointestinal infection. Med. 2022 Jun 10;3(6):371-387.e9.

8. Davidson MB, Peters AL. An overview of metformin in the treatment of type 2 diabetes mellitus. Vol. 102, American Journal of Medicine. 1997. p. 99–110.

9. Zhou G, Myers R, Li Y, Chen Y, Shen X, Fenyk-Melody J, et al. Role of AMP-activated protein kinase in mechanism of metformin action. J Clin Invest [Internet]. 2001 Oct 15 ;108(8):1167–74. Available from: http://www.jci.org.

10. Herzig S, Shaw RJ. AMPK: Guardian of metabolism and mitochondrial homeostasis. Nat Rev Mol Cell Biol [Internet]. 2018;19(2):121–35. Available from: http://dx.doi.org/10.1038/nrm.2017.95

11. Kim J, Yang G, Kim Y, Kim J, Ha J. AMPK activators: mechanisms of action and physiological activities. Exp Mol Med 2016 484 [Internet]. 2016 Apr 1;48(4):e224–e224. Available from: https://www.nature.com/articles/emm201616

12. Bramante CT, Ingraham NE, Murray TA, Marmor S, Hovertsen S, Gronski J, et al. Metformin and risk of mortality in patients hospitalised with COVID-19: a retrospective cohort analysis. Lancet Heal Longev [Internet]. 2021;2(1):e34– 41. Available from: http://dx.doi.org/10.1016/S2666-7568(20)30033-7

13. Dardano A, Del Prato S. Metformin: an inexpensive and effective treatment in people with diabetes and COVID-19? Lancet Heal Longev [Internet]. 2021;2(1):e6–7. Available from: http://dx.doi.org/10.1016/S2666-7568(20)30047-7

14. Ibrahim S, Lowe JR, Bramante CT, Shah S, Klatt NR, Sherwood N, et al. Metformin and Covid-19: Focused Review of Mechanisms and Current Literature Suggesting Benefit. Front Endocrinol (Lausanne). 2021 Jul 22;0:625.

15. Zangiabadian M, Nejadghaderi SA, Zahmatkesh MM, Hajikhani B, Mirsaeidi M, Nasiri MJ. The Efficacy and Potential Mechanisms of Metformin in the Treatment of COVID-19 in the Diabetics: A Systematic Review. Front Endocrinol (Lausanne). 2021;12(March):1–9.

16. L Z, ZG S, X C, JJ Q, XJ Z, J C, et al. Association of Blood Glucose Control and Outcomes in Patients with COVID-19 and Pre-existing Type 2 Diabetes. Cell Metab [Internet]. 2020 Jun 2;31(6):1068-1077.e3. Available from: https://pubmed.ncbi.nlm.nih.gov/32369736/

17. Marmor S, Bramante CT, Ingraham NE, Murray TA, Marmor S, Hovertsen S, et al. Metformin and risk of mortality in patients hospitalised with COVID-19: a retrospective cohort analysis. Artic Lancet Heal Longev [Internet]. 2021;2:34– 41. Available from: http://www.thelancet.com/

18. Chen X, Guo H, Qiu L, Zhang C, Deng Q, Leng Q. Immunomodulatory and Antiviral Activity of Metformin and Its Potential Implications in Treating Coronavirus Disease 2019 and Lung Injury. Front Immunol. 2020 Aug 18;0:2056.

19. Chen Y, Chen L, Deng Q, Zhang G, Wu K, Ni L, et al. The presence of SARS-CoV-2 RNA in the feces of COVID-19 patients. J Med Virol [Internet]. 2020 Jul 1;92(7):833–40. Available from: https://pubmed.ncbi.nlm.nih.gov/32243607/

20. Lamers MM, Beumer J, Vaart J Van Der, Knoops K, Puschhof J, Breugem TI, et al. SARS-CoV-2 productively infects human gut enterocytes. Science (80-) [Internet]. 2020 Jul 3;369(6499):50–4. Available from: https://www.science.org/doi/10.1126/science.abc1669

21. Gupta D, Parthasarathy H, Sah V, Tandel D, Vedagiri D, Reddy S, et al. Inactivation of SARS-CoV-2 by β-propiolactone causes aggregation of viral particles and loss of antigenic potential. Virus Res [Internet]. 2021 Nov 1;305:198555. Available from: https://linkinghub.elsevier.com/retrieve/pii/S0168170221002628

22. Schneider CA, Rasband WS, Eliceiri KW. NIH Image to ImageJ: 25 years of image analysis. Nat Methods 2012 97 [Internet]. 2012 Jun 28;9(7):671–5. Available from: https://www.nature.com/articles/nmeth.2089

23. Bhutta MS, Gallo ES, Borenstein R. Multifaceted Role of AMPK in Viral Infections. Cells [Internet]. 2021;10(5). Available from: /pmc/articles/PMC8148118/

24. Tsai H-H, Lai H-Y, Chen Y-C, Li C-F, Huang H-S, Liu H-S, et al. Metformin promotes apoptosis in hepatocellular carcinoma through the CEBPD-induced autophagy pathway. Oncotarget [Internet]. 2017 Jan 13;8(8):13832–45. Available from: https://www.oncotarget.com/article/14640/text/

25. Cauchy F, Mebarki M, Leporq B, Laouirem S, Albuquerque M, Lambert S, et al. Strong antineoplastic effects of metformin in preclinical models of liver carcinogenesis. Clin Sci (Lond) [Internet]. 2017;131(1):27–36. Available from: https://pubmed.ncbi.nlm.nih.gov/27803295/

26. Alhourani AH, Tidwell TR, Bokil AA, Røsland G V., Tronstad KJ, Søreide K, et al. Metformin treatment response is dependent on glucose growth conditions and metabolic phenotype in colorectal cancer cells. Sci Reports 2021 111 [Internet]. 2021 May 18;11(1):1–10. Available from: https://www.nature.com/articles/s41598-021-89861-6

27. Sanchez-Rangel E, Inzucchi SE. Metformin: clinical use in type 2 diabetes. Diabetologia [Internet]. 2017 Sep 1;60(9):1586–93. Available from: https://link.springer.com/article/10.1007/s00125-017-4336-x

28. Fernandes-Siqueira LO, Zeidler JD, Sousa BG, Ferreira T, Da Poian AT. Anaplerotic Role of Glucose in the Oxidation of Endogenous Fatty Acids during Dengue Virus Infection. mSphere [Internet]. 2018 Feb 28;3(1). Available from: https://journals.asm.org/doi/abs/10.1128/mSphere.00458-17

29. Shen C, Ka SO, Kim SJ, Kim JH, Park BH, Park JH. Metformin and AICAR regulate NANOG expression via the JNK pathway in HepG2 cells independently of AMPK. Tumor Biol 2016 378 [Internet]. 2016 Mar 3;37(8):11199–208. Available from: https://link.springer.com/article/10.1007/s13277-016-5007-0

30. Quan HY, Kim DY, Chung SH. Caffeine attenuates lipid accumulation via activation of AMP-activated protein kinase signaling pathway in HepG2 cells. BMB Rep [Internet]. 2013;46(4):207–12. Available from: http://dx.

31. Huang C, Wang Y, Li X, Ren L, Zhao J, Hu Y, et al. Clinical features of patients infected with 2019 novel coronavirus in Wuhan, China. Lancet [Internet]. 2020 Feb 15;395(10223):497–506. Available from: http://www.thelancet.com/article/S0140673620301835/fulltext

32. Crouse AB, Grimes T, Li P, Might M, Ovalle F, Shalev A. METFORMIN USE IS ASSOCIATED WITH REDUCED MORTALITY IN A DIVERSE POPULATION WITH COVID-19 AND DIABETES. medRxiv [Internet]. 2020 Jul 31;2020.07.29.20164020. Available from: https://www.medrxiv.org/content/10.1101/2020.07.29.20164020v1

33. P L, L Q, Y L, XL L, JL Z, HY X, et al. Metformin Treatment Was Associated with Decreased Mortality in COVID-19 Patients with Diabetes in a Retrospective Analysis. Am J Trop Med Hyg [Internet]. 2020 Jul 1;103(1):69– 72. Available from: https://pubmed.ncbi.nlm.nih.gov/32446312/

34. Lui DTW, Tan KCB. Is metformin a miracle or a menace in COVID-19 patients with type 2 diabetes? J Diabetes Investig [Internet]. 2021 Apr 1;12(4):479–81. Available from: https://onlinelibrary.wiley.com/doi/full/10.1111/jdi.13484

35. Pernicova I, Korbonits M. Metformin—mode of action and clinical implications for diabetes and cancer. Nat Rev Endocrinol 2014 103 [Internet]. 2014 Jan 7;10(3):143–56. Available from: https://www.nature.com/articles/nrendo.2013.256

36. Rena G, Hardie DG, Pearson ER. The mechanisms of action of metformin. Diabetologia. 2017;60(9):1577–85.

37. Pereira-Dutra FS, Teixeira L, Costa MF de S, Bozza PT. Fat, fight, and beyond: The multiple roles of lipid droplets in infections and inflammation. J Leukoc Biol [Internet]. 2019 Sep 1;106(3):563–80. Available from: https://onlinelibrary.wiley.com/doi/full/10.1002/JLB.4MR0119-035R

38. Herker E, Ott M. Emerging Role of Lipid Droplets in Host/Pathogen Interactions. J Biol Chem. 2012 Jan 1;287(4):2280–7.

39. Syed GH, Siddiqui A. Effects of hypolipidemic agent nordihydroguaiaretic acid on lipid droplets and Hepatitis C virus. Hepatology [Internet]. 2011 Dec;54(6):1936. Available from: /pmc/articles/PMC3236615/

40. Széles L, Töröcsik D, Nagy L. PPARγ in immunity and inflammation: cell types and diseases. Biochim Biophys Acta - Mol Cell Biol Lipids. 2007 Aug 1;1771(8):1014–30.

41. MM S, JA M, NG I, I A-M, G B-L, AT DP, et al. Dengue virus capsid protein usurps lipid droplets for viral particle formation. PLoS Pathog [Internet]. 2009 Oct;5(10). Available from: https://pubmed.ncbi.nlm.nih.gov/19851456/

42. Y M, K A, N U, K W, T H, M Z, et al. The lipid droplet is an important organelle for hepatitis C virus production. Nat Cell Biol [Internet]. 2007 Sep;9(9):1089– 97. Available from: https://pubmed.ncbi.nlm.nih.gov/17721513/

43. PS S, N L, GM J, PP S, T Z, DL S, et al. Comparative Flavivirus-Host Protein Interaction Mapping Reveals Mechanisms of Dengue and Zika Virus Pathogenesis. Cell [Internet]. 2018 Dec 13;175(7):1931-1945.e18. Available from: https://pubmed.ncbi.nlm.nih.gov/30550790/

44. S B, P T-A, J M. Disrupting the association of hepatitis C virus core protein with lipid droplets correlates with a loss in production of infectious virus. J Gen Virol [Internet]. 2007 Aug;88(Pt 8):2204–13. Available from: https://pubmed.ncbi.nlm.nih.gov/17622624/

45. Doerflinger SY, Cortese M, Romero-Brey I, Menne Z, Tubiana T, Schenk C, et al. Membrane alterations induced by nonstructural proteins of human norovirus. PLOS Pathog [Internet]. 2017 Oct 1;13(10):e1006705. Available from: https://journals.plos.org/plospathogens/article?id=10.1371/journal.ppat.1006705

46. Coffey CM, Sheh A, Kim IS, Chandran K, Nibert ML, Parker JSL. Reovirus Outer Capsid Protein μ1 Induces Apoptosis and Associates with Lipid Droplets, Endoplasmic Reticulum, and Mitochondria. J Virol [Internet]. 2006 Sep;80(17):8422. Available from: /pmc/articles/PMC1563861/

47. W C, M G, A E, CF K, N C, S C, et al. Rotaviruses associate with cellular lipid droplet components to replicate in viroplasms, and compounds disrupting or blocking lipid droplets inhibit viroplasm formation and viral replication. J Virol [Internet]. 2010 Jul;84(13):6782–98. Available from: https://pubmed.ncbi.nlm.nih.gov/20335253/

48. Monson EA, Crosse KM, Das M, Helbig KJ. Lipid droplet density alters the early innate immune response to viral infection. PLoS One [Internet]. 2018 Jan 1;13(1):e0190597. Available from: https://journals.plos.org/plosone/article?id=10.1371/journal.pone.0190597

49. Dias SSG, Soares VC, Ferreira AC, Sacramento CQ, Fintelman-Rodrigues N, Temerozo JR, et al. Lipid droplets fuel SARS-CoV-2 replication and production of inflammatory mediators. PLOS Pathog [Internet]. 2020 Dec 16;16(12):e1009127. Available from: https://journals.plos.org/plospathogens/article?id=10.1371/journal.ppat.1009127

50. Proal AD, VanElzakker MB. Long COVID or Post-acute Sequelae of COVID-19 (PASC): An Overview of Biological Factors That May Contribute to Persistent Symptoms. Front Microbiol. 2021 Jun 23;12:1494.

51. Chu H, Chan JF-W, Yuen TT-T, Shuai H, Yuan S, Wang Y, et al. Comparative tropism, replication kinetics, and cell damage profiling of SARS-CoV-2 and SARS-CoV with implications for clinical manifestations, transmissibility, and laboratory studies of COVID-19: an observational study. The Lancet Microbe [Internet]. 2020 May;1(1):e14–23. Available from: https://pubmed.ncbi.nlm.nih.gov/32835326/

52. Hoffmann M, Kleine-Weber H, Schroeder S, Krüger N, Herrler T, Erichsen S, et al. SARS-CoV-2 Cell Entry Depends on ACE2 and TMPRSS2 and Is Blocked by a Clinically Proven Protease Inhibitor. Cell. 2020;181(2):271-280.e8.

53. Foretz M, Guigas B, Bertrand L, Pollak M, Viollet B. Metformin: From Mechanisms of Action to Therapies. Cell Metab [Internet]. 2014 Dec 2;20(6):953–66. Available from: http://www.cell.com/article/S1550413114004410/fulltext

54. Bojkova D, Klann K, Koch B, Widera M, Krause D, Ciesek S, et al. Proteomics of SARS-CoV-2-infected host cells reveals therapy targets. Nature [Internet]. 2020;583(7816):469–72. Available from: http://dx.doi.org/10.1038/s41586-020-2332-7

